# Prospects and limitations of expansion microscopy in chromatin ultrastructure determination

**DOI:** 10.1101/2020.03.16.994186

**Authors:** Ivona Kubalová, Markéta Schmidt Černohorská, Martina Huranová, Klaus Weisshart, Andreas Houben, Veit Schubert

**Affiliations:** Leibniz Institute of Plant Genetics and Crop Plant Research (IPK) Gatersleben, D-06466 Seeland, Germany; Laboratory of Adaptive Immunity, Institute of Molecular Genetics, Academy of Sciences of the Czech Republic, Prague, Czech Republic; Carl Zeiss Microscopy GmbH, D-07745 Jena, Germany

**Keywords:** chromatin, expansion microscopy, nucleus, structured illumination microscopy, *Hordeum vulgare*

## Abstract

Expansion Microscopy (ExM) is a method to magnify physically a specimen with preserved ultrastructure. It has the potential to explore structural features beyond the diffraction limit of light. The procedure has been successfully used for different animal species, from isolated macromolecular complexes through cells to tissue slices. Expansion of plant-derived samples is still at the beginning, and little is known whether the chromatin ultrastructure becomes altered by physical expansion.

In this study, we expanded isolated barley nuclei and compared whether ExM can provide a structural view of chromatin comparable with super-resolution microscopy. Different fixation and denaturation/digestion conditions were tested to maintain the chromatin ultrastructure. We achieved up to ∼4.2-times physically expanded nuclei corresponding to a maximal resolution of ∼50-60 nm when imaged by wild-field (WF) microscopy. By applying structured illumination microscopy (SIM, super-resolution) doubling the WF resolution the chromatin structures were observed at a resolution of ∼25-35 nm.

WF microscopy showed a preserved nucleus shape and nucleoli. Moreover, we were able to detect chromatin domains, invisible in unexpanded nuclei. However, by applying SIM we observed that the preservation of the chromatin ultrastructure after expansion was not complete and that the majority of the tested conditions failed to keep the ultrastructure.

Nevertheless, using expanded nuclei we detected successfully centromere repeats by fluorescence *in situ* hybridization (FISH) and the centromere-specific histone H3 variant CENH3 by indirect immunostaining. However, although these repeats and proteins were localized at the correct position within the nuclei (indicating a Rabl orientation) their ultrastructural arrangement was impaired.

## Introduction

Expansion microscopy (ExM) is a method to enlarge small structures physically in an isotropic manner to overcome the diffraction limit of light microscopy. Thus, super-resolution (<250 nm) can be realized cost-efficiently with diffraction-limited light microscopes (**Chen et al. 2015**; **Chang et al. 2017**). Even a lateral resolution of ∼70 nm can be achieved by combining ExM and standard confocal microscopy (**Jiang et al. 2018)**.

ExM is based on a swellable polyelectrolyte gel, increasing in size when exposed to water to achieve a ∼4.5-fold 3-dimensional (3D) expansion (**Alon et al. 2019; Wassie et al. 2019**). The first ExM protocol expanding mouse brain tissue 4.5-times was described by **Chen et al. (2015**). Since then, several expansion protocols emerged to increase the expansion factor and to preserve the ultrastructural features. These protocols were adapted to species like fungi, human, mouse, fruit fly, zebrafish and soft tissues such as brain, skin, kidney and liver (**Chen et al. 2015; Tillberg et al., 2016; Cahoon et al. 2017; Freifeld et al. 2017; Halpern et al. 2017; Jiang et al. 2018, Lim et al. 2019; Truckenbrodt et al., 2019; Zwettler et al. 2019; Götz et al. 2020**). The following processes occur during ExM to fix, embed and expand the specimen successfully: 1) during the fixation with a formaldehyde/acrylamide mixture formaldehyde crosslinks proteins/DNA/RNA to each other; 2) during gelation the crosslinked proteins become crosslinked to the polyacrylamide (PAA) gel due to the acrylamide provided during fixation; 3) during denaturation in SDS buffer and at high temperature, all crosslinked proteins denature while remaining crosslinked to the PAA gel mesh which starts to expand in the denaturation buffer; 4) during expansion in water, all proteins renature back with gaps between each other, but still bound to the PAA gel mesh preserving their exact position as before expansion (**Chen et al. 2015; Cho et al. 2018; Tillberg and Chen 2019; Wassie et al. 2019**).

At the subcellular level, expansion and super-resolution microscopy have been combined to analyse fruit fly and mouse synaptonemal complex protein components (**Cahoon et al. 2017; Wang et al. 2018b; Xu et al. 2019)**. Super-resolution microscopy techniques such as structured illumination microscopy (SIM) are subdiffraction imaging methods bridging the resolution gap between light and electron microscopy. They were applied successfully in cell biology (**Fornasiero et al. 2015**) at specimens from both prokaryotes and eukaryotes and allowed also discovering new structures within plant chromatin (**Schubert 2017**). The multiplication of the achieved physical and optical resolution of both methods could also be useful to decipher the 3D structure of chromatin in cell nuclei and highly condensed metaphase chromosomes.

ExM was successfully applied to visualize specific proteins and RNAs by immunostaining and *in situ* hybridization, respectively (**Chen et al. 2016; Chozinski et al. 2016; Asano et al. 2018**). Labelling of specific DNA sequences in spatially expanded chromatin has not yet been reported. Only the application of DNA-specific dye like DAPI was shown in combination with ExM (**Zhao et al. 2017; Düring et al. 2019)**.

Physically expanded nuclei and chromosomes in combination with optical super-resolution microscopy to increase the resolution would allow analysing ultrastructure, dynamics and function of chromatin more in detail, especially via the detection of DNA sequences and proteins after specific fluorescence labelling. Until now the preservation of the ultrastructure of expanded chromatin has not yet been analysed by super-resolution microscopy. A previous study showed that isolated barley chromosomes can become expanded after gentle fixation and flow-sorting (**Endo et al. 2014**). However, whether the chromatin ultrastructure of these expanded chromosomes is preserved has not been analysed.

Caused by varying refractive indices of plant cell organelles, which induce spherical aberrations and light scattering (**Komis et al. 2015**), plant cell imaging is more challenging than imaging of prokaryotic and animal/human tissues. Due to the absence of cytoplasm, isolated and flow-sorted nuclei are well suitable to perform immunostaining and FISH followed by SIM (**Schubert and Weisshart 2015; Weisshart et al. 2016; Schubert 2017**).

To test whether expansion microscopy could be applied to improve the ultrastructural analysis of somatic plant chromatin, we isolated interphase nuclei of barley and tested different preparation methods based on an advanced ExM protocol for ultrastructures, called ultra-expansion microscopy (U-ExM) (**Gambarotto et al. 2019**). We achieved a physical ∼4.2-fold nuclei expansion and the partial preservation of the chromatin ultrastructure as proven by standard wide-field microscopy. Besides, ExM was combined with immunolabelling and FISH to analyse the interphase centromeres of barley. However, after analysis of expanded chromatin by super-resolution microscopy, we noticed that the chromatin substructure was altered due to ExM.

## Materials and Methods

### Plant material and nuclei isolation

Root tips of barley (*Hordeum vulgare* L. var. “Morex”) seedlings were collected in a fixation solution (formaldehyde (FA), glutaraldehyde (GA) or glyoxal) and treated 5 min under vacuum, followed by incubation on ice for the indicated time (Suppl. Table 1). After fixation, at least 100 root tips were washed twice with a 1×PBS solution and immediately chopped using a razor blade in 400 µl nuclei-isolation buffer LB01 (**Doležel et al. 2007**). The nuclei suspension was filtered using a 50 µm filter mesh (CellTrics^®^, SYSMEX), collected into a new tube and stained with 4′,6-diamidino-2-phenylindole (DAPI) (∼5 µg/ml, Molecular Probes # D1306). Round coverslips (Ø12 mm) (Thermo Scientific, Menzel Gläser) were placed into a 24-well culture plate (Greiner) and coated with poly-L-lysine for at least 20 min. The dispensable poly-L-lysine solution was removed and the prepared coverslips were used immediately or stored at room temperature (RT) in wells for later use. To load a coverslip with nuclei the nuclei suspension was pipetted into the well containing a coverslip and centrifuged at 1,000 g for 10 min at RT using a swing-bucket rotor (Eppendorf Centrifuge 5810 R). The supernatant can be re-used for additional coverslips.

### Gelation, denaturation/digestion and expansion

Before gelation, the nuclei-loaded coverslip was taken out from the 24-well culture well without drying. Per gel 50 µl monomer solution (MS) (Suppl. Table 2) was mixed with tetramethylethylenediamine (TEMED) and ammonium persulfate (APS) (final concentration 0.2% w/w), and 35 µl of the resulting mixture was dropped onto the clean surface of an ice-cold plastic plate covered by parafilm. The nuclei-loaded coverslip was promptly placed on top of the gel drop with nuclei facing the gel. The plate was kept on ice for 5 min to allow the gel to solidify. To finalize the solidification process, the gel was placed into a wet chamber and incubated for 15 min at 37°C. The solidified gel was carefully removed from the coverslip using flat forceps and submerged into either a denaturation or digestion buffer (Suppl. Table 2) or water, and incubated as designed in Suppl. Table 1. The simplified schemes of the protocol steps are shown for monomer solution 1 and 2 in Figures 1a and 1b, respectively. In the case of a digestion step, before the incubation with a digestion buffer, proteinase K was added at a final concentration of 8 U/ml. After denaturation/digestion the gel was expanded in distilled water until the expansion reached the maximum possible size of ∼50.4 mm (∼4.2 times expansion). The distilled water was changed at least three times. Due to the presence of the nuclei population from distinct cell cycle stages and technical impossibility to visualize nuclei in the gel before expansion, the expansion factor was estimated only on the gel expansion, from 12 to 50.4 mm.

**Figure 1:**
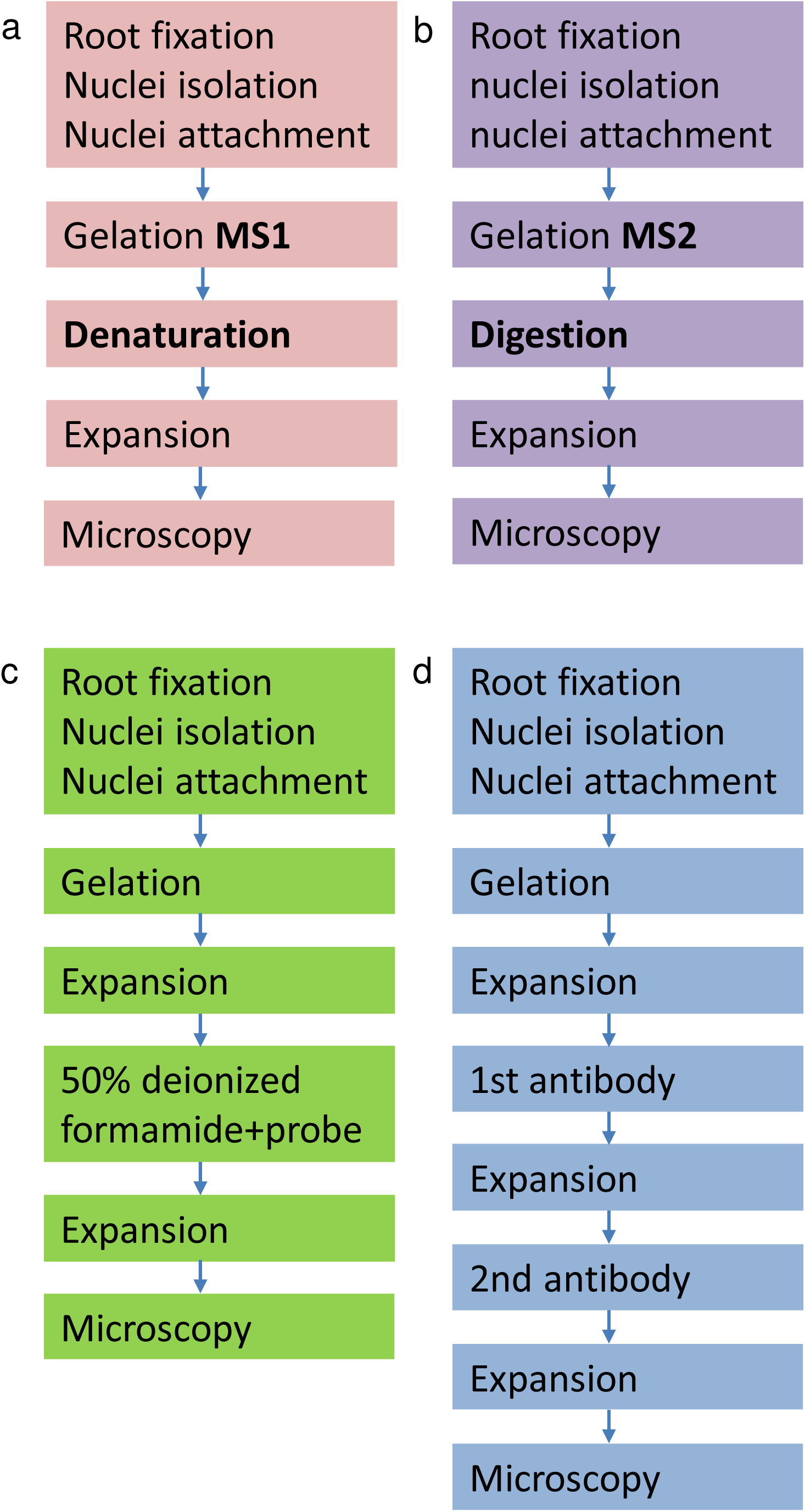
Simplified schemes of ExM protocols for isolated nuclei. The standard ExM protocols for monomer solution 1 and 2 are shown in (a) and (b), respectively. The protocols were optimised for FISH (c) and indirect immunolabelling (d). The denaturation/digestion step was excluded from the optimised protocols (c, d).

### Non-denaturating fluorescence *in situ* hybridization (ND-FISH)

A nuclei-containing fragment of the expanded gel was incubated in 50% deionized formamide in 2×SSC for 20 min at RT. Afterwards, formamide was replaced by 50 µl of hybridization mixture containing 50% deionized formamide, 2×SSC and 2 µg of the 5’FAM-labelled barley centromere-specific oligo probe (GA)_15_. The gel was incubated in a sealed chamber for 22 h at 37°C. The exposure to the hybridization solution containing ions (hypertonic) results in shrinkage of the expanded gel. Therefore, after hybridization, the remaining hybridization mixture was removed and the gel was re-expanded in distilled water followed by microscopical observation. The simplified scheme of these protocol steps is shown in Fig. 1c.

### Indirect immunostaining

A nuclei-containing fragment of the expanded gel was incubated with primary rabbit antibodies against the centromeric H3 variant CENH3 of barley (**Houben et al. 2007**) diluted 1:1000 in 400 µl of antibody solution (2.5% BSA, 0.05% Triton X-100, 1×PBS) in a 12-well culture plate (Greiner) and incubated overnight for at least 15 h at RT. Next, the primary antibody solution was removed and the gel was re-expanded in distilled water until it reached the size before antibody incubation. Afterwards, the gel was incubated with secondary anti-rabbit Alexa488 antibodies (1:200, #711-545-152 Jackson ImmunoResearch) in 400 µl antibody solution and incubated at 37°C for 1 h followed by 2 h at RT. Additionally, 3-5 µg/ml DAPI solution can be added together with the secondary antibodies to enable chromatin staining. Finally, the gel was re-expanded in distilled water and subjected to microscopic observation in a 22×22 mm coverslip chamber ‘Chamlide’ (Live Cell Instruments, catalogue no. CM-S22-1). The simplified scheme of these protocol steps is shown in Fig. 1d.

### Widefield, deconvolution and super-resolution microscopy

The chromatin structure was analysed by widefield (WF), deconvolution (DCV) of WF and super-resolution, using an Elyra PS.1 microscope system and the software ZEN Black (Carl Zeiss GmbH). Images were captured separately for DAPI and Alexa488 using the 405 nm and 488 laser lines for excitation and appropriate emission filters. To analyse the chromatin ultrastructure at a resolution of ∼100 nm (super-resolution achieved with a 405 nm laser) structured illumination microscopy (SIM) was performed with a 63×/1.4 Oil Plan-Apochromat objective (**Weisshart et al. 2016**).

For SIM imaging of unexpanded and expanded nuclei, a linear grid matching to the respective wavelength was used and the raw data were processed using the SIM processing function of ZEN Black. First, we started with the automatic mode and then optimized systematically to the highest strength of the noise filter where structured noise just disappeared. The resolution in the images was measured with the profile tool of ZEN Black taking a peak to peak distances between two structures. In theory, the resolution in expanded samples could be as high as SIM resolution divided by the expansion factor, assuming that the sample expansion was isotropic. The procedure to process the SIM raw data and to estimate the achieved resolution of unexpanded and expanded nuclei after WF and SIM imaging is described in detail in **Kubalová, et al. (2020**).

The WF and deconvoluted WF images were calculated in parallel to SIM processing by ZEN Black.

## Results

### A ∼4.2-times expansion of isolated plant nuclei can be achieved without denaturation and digestion

To establish a protocol to expand isolated interphase nuclei while preserving the chromatin ultrastructure several fixation conditions and two different monomer solution (MS) compositions were tested for expansion microscopy (ExM) (Suppl. Table 1). To analyse the effect of expansion on the chromatin ultrastructure we applied wide-field (WF), deconvolution (DCV) of WF images and structured illumination microscopy (SIM). Unexpanded nuclei were imaged by all three techniques as the untreated control to evaluate whether the imaging of expanded nuclei by WF could deliver structural information comparable to DCV or SIM applied to unexpanded nuclei (Fig. 2). Additionally, SIM on expanded nuclei was applied to assess whether the chromatin ultrastructure was altered due to the swelling of nuclei (Figs. 3-6).

**Figure 2:**
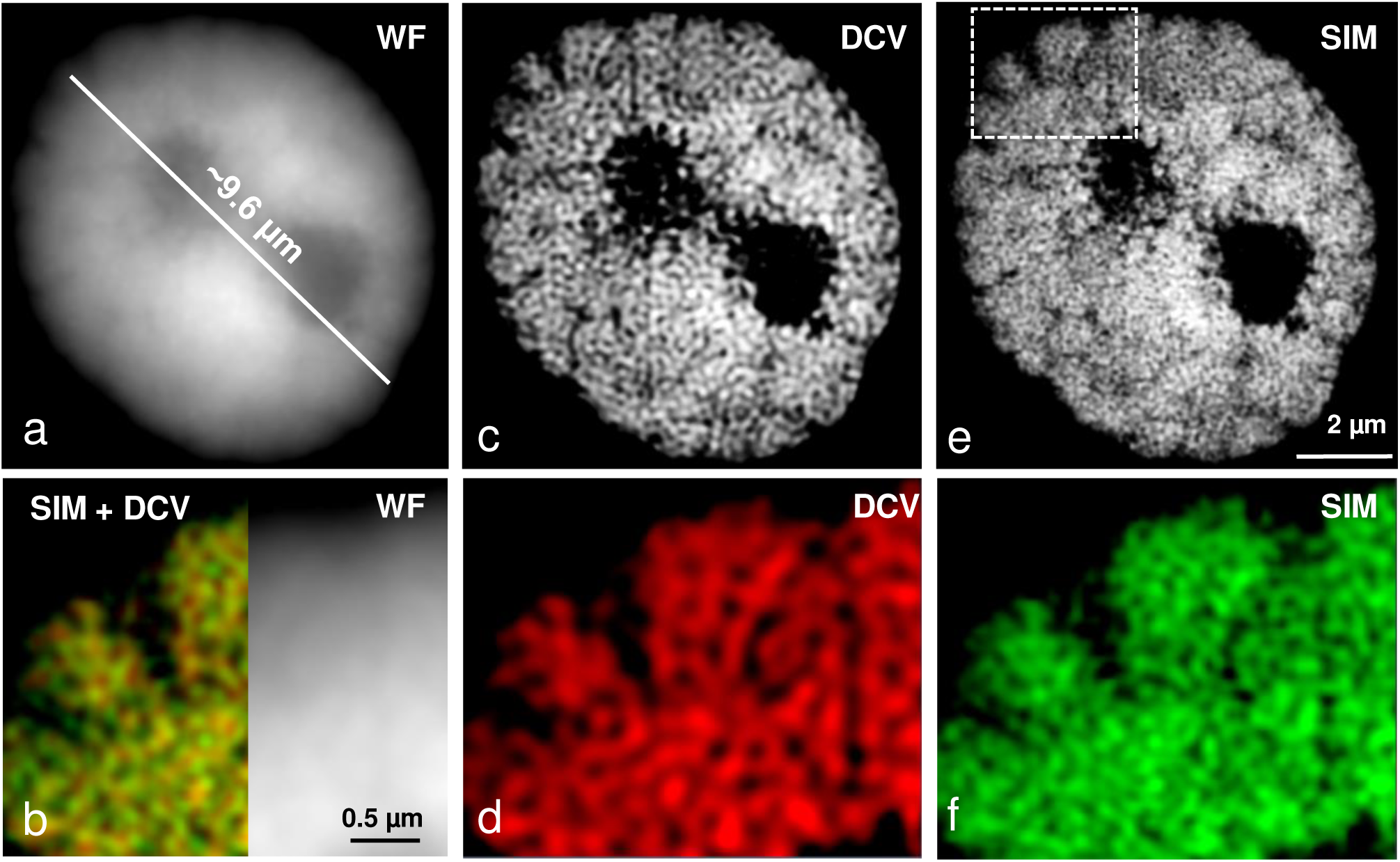
The chromatin fine structure is well maintained in an unexpanded nucleus, as especially visible after structured illumination microscopy (SIM) compared to deconvolution (DCV) and widefield microscopy (WF). Despite the different achieved resolution (a, c, e) the merge (b, left) of DCV (c) and SIM (e) indicates in the enlarged region (dashed rectangle) (b, d, f) that the same chromatin structures were identified. Instead, no clear structures are visible in the zoomed region by WF (b, right). Global chromatin was labelled by DAPI.

**Figure 3:**
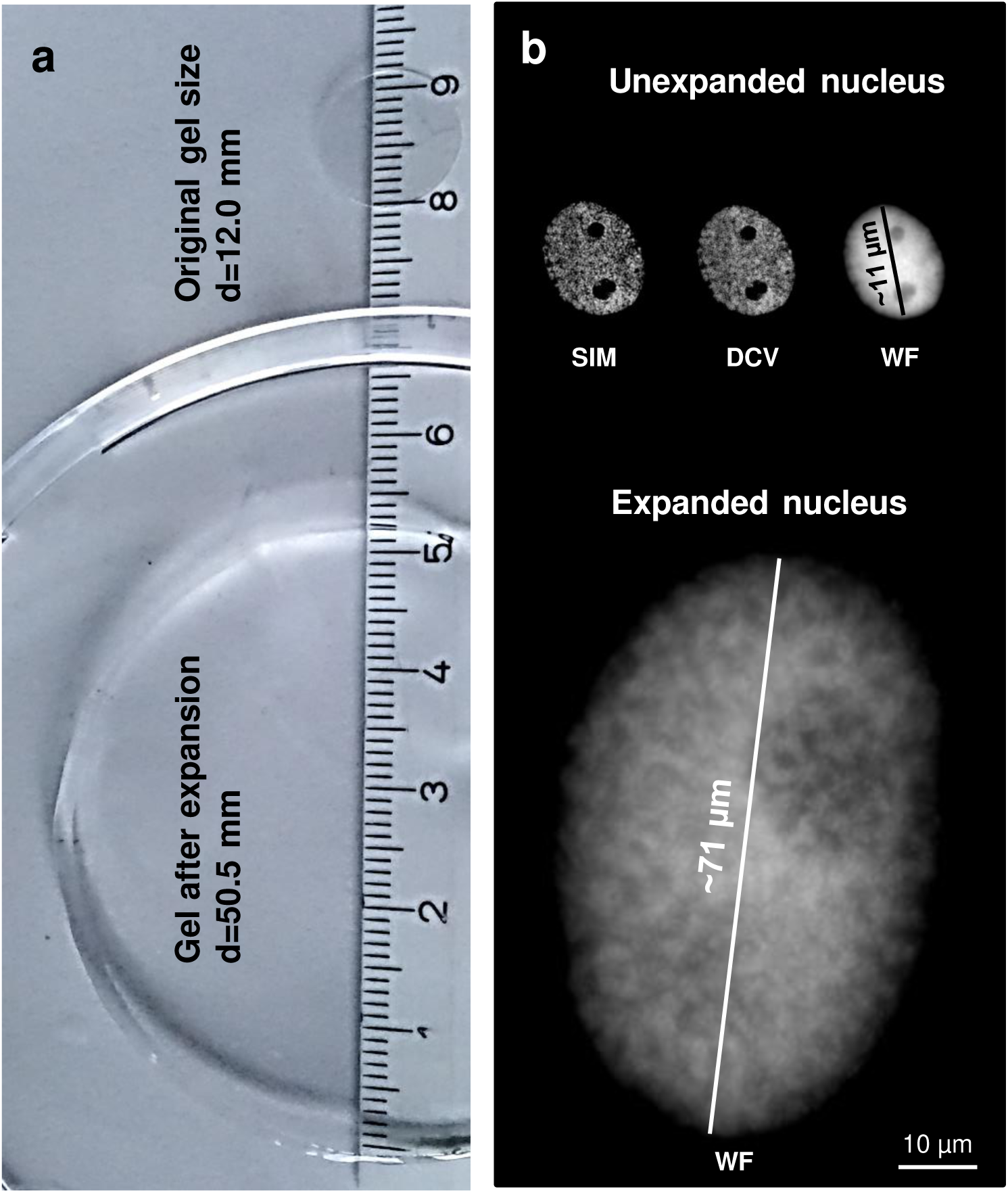
The expansion factor of gel (a) and nuclei (b) correspond to ∼4.2. Nuclei were stained by DAPI and imaged by structured illumination microscopy (SIM), deconvolution (DCV) and widefield microscopy (WF). Imaged by WF the expanded nucleus shows more structures than the original one, but does not reach the resolution of SIM.

**Figure 4:**
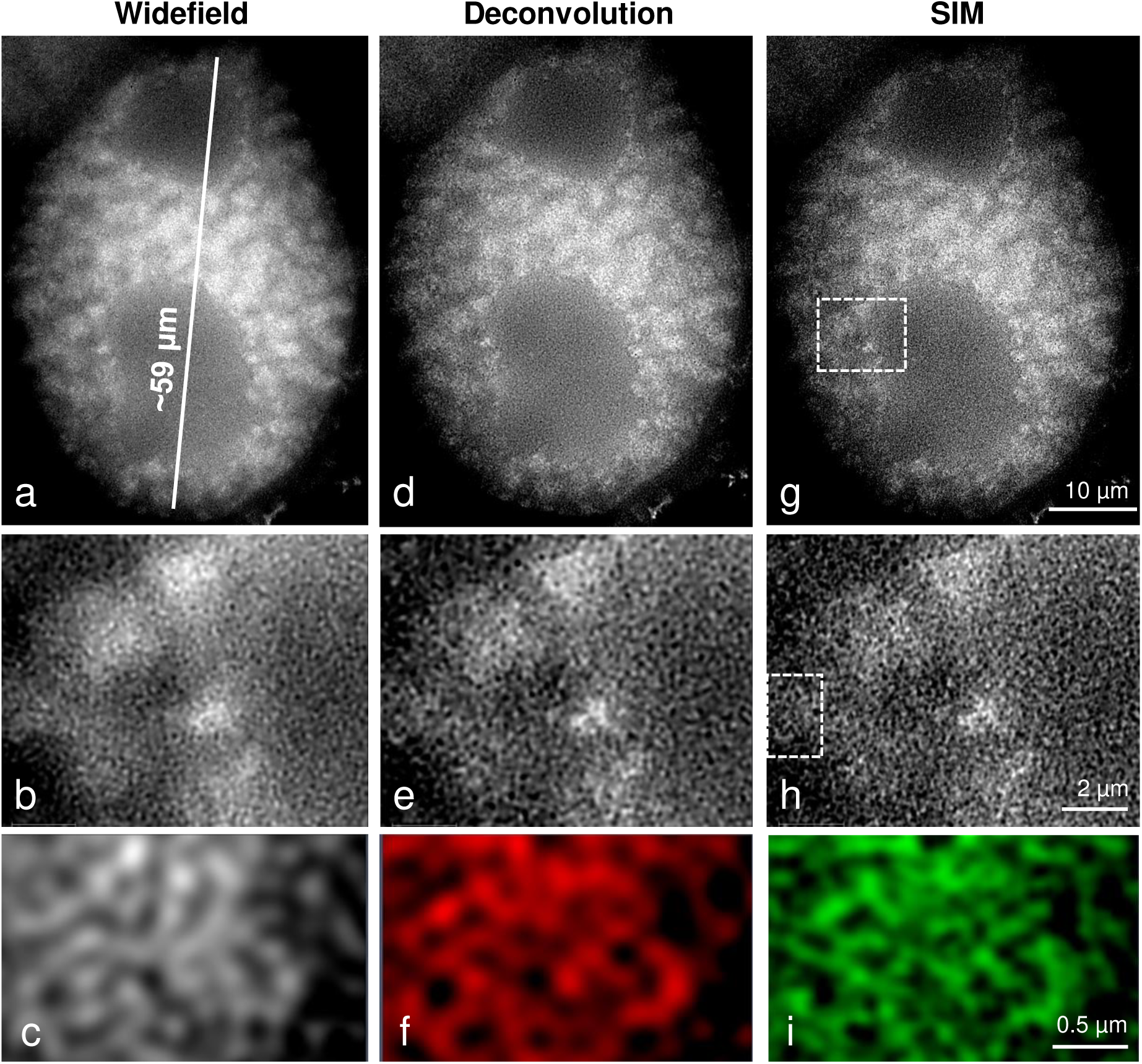
Well preserved chromatin structure achieved by application of ExM protocol variant 21 (see suppl. table 1). The subsequent magnifications of selected regions (dashed rectangles) demonstrate the preserved chromatin fine structure visible even after wide-field microscopy (a, b, c) of this completely expanded nucleus labelled by DAPI. By comparing WF with deconvolution (d, e, f) and SIM (g, h, i) similar chromatin structures become visible at the highest magnification (region 90° rotated).

**Figure 5:**
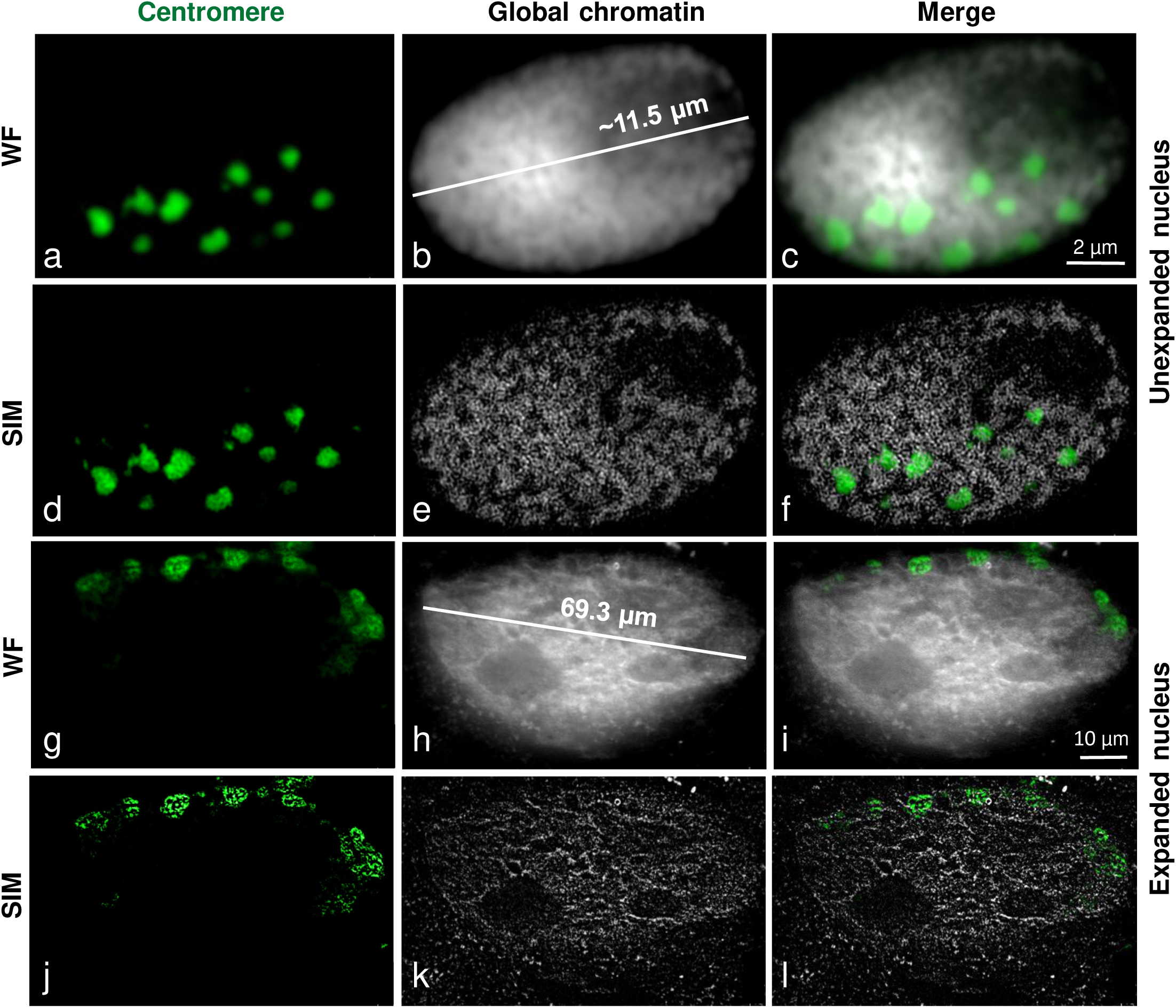
FISH detection of centromeric repeats (GA)15 at unexpanded (a-f) and expanded (g-l) nuclei using ExM protocol variant 21 (see suppl. table 1). Although the main centromeric structures are maintained after complete expansion and thus indicating Rabl orientation WF (g, h, i) and SIM (j, k, l) imaging show that the chromatin ultrastructures are impaired. Global chromatin was labelled by DAPI.

**Figure 6:**
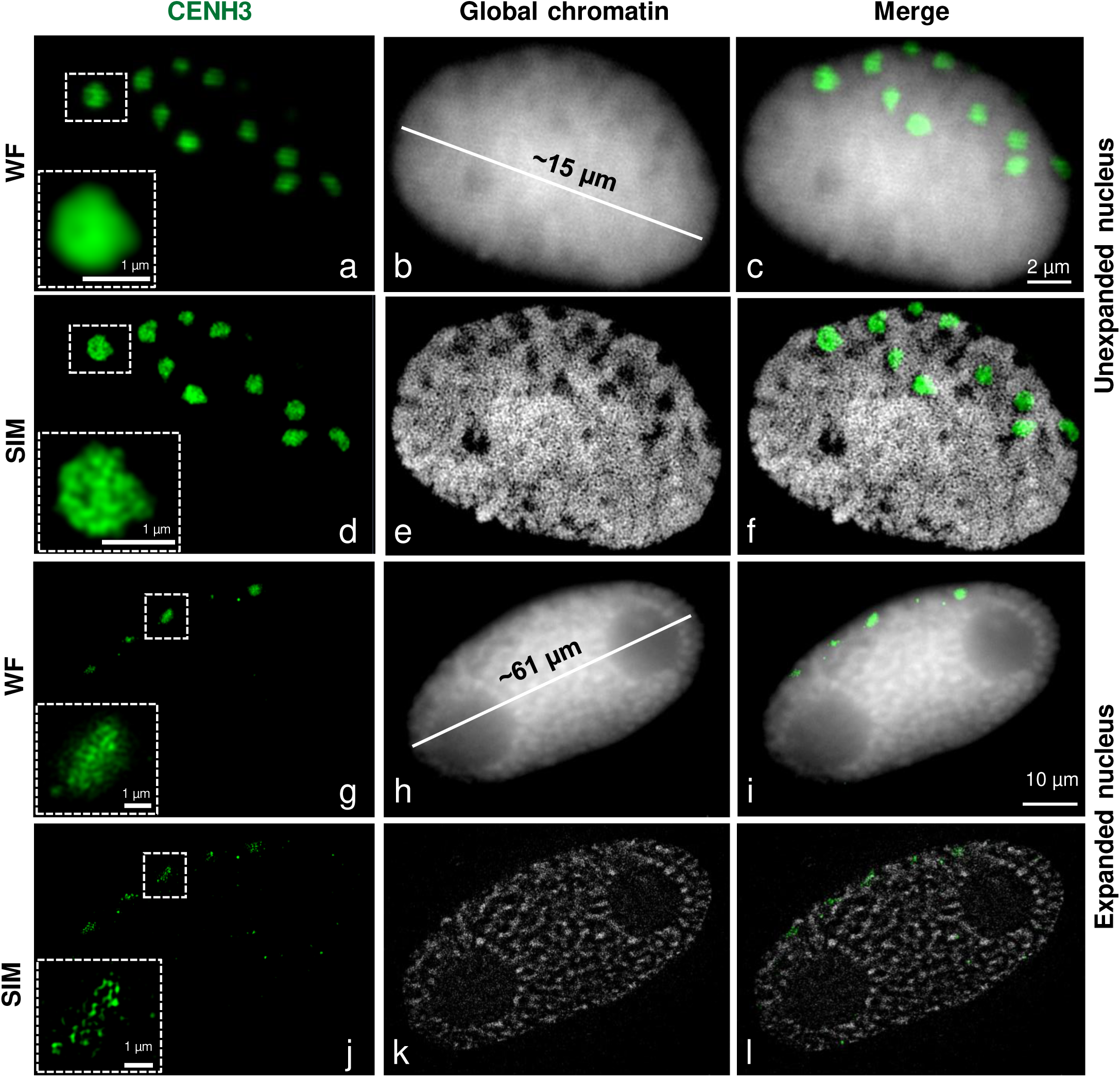
Detection of the centromere-specific histone variant CENH3 after immunolabeling and ExM according to protocol variant 6 (see suppl. table 1). Although the main centromeric structures are preserved after complete expansion, SIM and WF imaging indicate that the chromatin fine structures are impaired. The enlarged region (dashed rectangle) shows clearly the preserved and distorted CENH3-positive chromatin in an unexpanded (a-f) and expanded (g-l) nucleus, respectively. Global chromatin was labelled by DAPI.

First, we performed 12 min fixation of barley root tips in a mixture of 2% formaldehyde (FA) + 2% acrylamide (AA) in 1×PBS prior the extraction of nuclei, combined with monomer solution MS1 with more concentrated AA (Suppl. Table 2), and applied heat denaturation for 3 min at 80°C in denaturation buffer containing SDS (Suppl. Table 2; protocol variant 2 in Suppl. Table 1). After expansion, we observed a ∼4.2-times expansion of the gel and nuclei. (Fig. 3; Suppl. Fig. 1). As the size of untreated nuclei varied between 9 to 25 µm in diameter due to the mixed tissue-type origin (tissue layers of the root tip) and status of DNA replication (G1, S and G2) also the size of expanded nuclei varied between ∼40 and ∼110 µm (Suppl. Fig. 2).

By using a 20×/0.8 objective we observed that all nuclei were expanded and showed similar structures (Suppl. Fig. 2). A few of them were selected for SIM analysis using the 63×/1.4 objective. Structural features, such as the nucleus shape and nucleoli, were always visible after expansion. Also, the chromatin arrangement into distinct domains representing chromatin associations became visible by WF (Suppl. Figs. 1a, b). However, the application of DCV and SIM revealed that the chromatin ultrastructure was impaired because chromatin fibres organized as a network and clearly visible in unexpanded nuclei by SIM (Fig. 2e, f), were not detectable (Suppl. Fig. 1e, f).

To improve the expansion protocol, we denatured the specimen for 3 min with 0.2 M NaOH in 70% ethanol, instead of heat denaturation. The alkaline/ethanol denaturation is commonly used to denature DNA during the FISH procedure and is superior for chromatin structure preservation (**Raap et al. 1986; Andras et al. 1999**). Again, we achieved a ∼4.2-times expansion of gel and nuclei. All microscopical imaging methods showed a better preserved chromatin ultrastructure of expanded nuclei. Even the application of WF microscopy showed structures like chromatin domains with a resolution of ∼50-60 nm that are invisible in unexpanded nuclei (Suppl. Figs. 3a-c; protocol variant 3 in Suppl. Table 1). To achieve an optimal SIM image quality the moderate noise filter setting of −3.8 was applied in the SIM calculation tool of the ZEN Black software. A filter setting of 1.0 representing a high strength for the calculation was only suitable for the calculation of unexpanded nuclei (**Weisshart et al. 2016**). Thus, expansion leads to a decrease in the signal-to-noise ratio (SNR) of the images, and hence to the reduction of the specific sample fluorescence. Nevertheless, in expanded nuclei, SIM imaging resulted in a resolution of ∼25-35 nm.

Next, to simplify the protocol and further improve the chromatin ultrastructure preservation we changed the fixation solution by using a combination of 1% FA and 1% glutaraldehyde and applied 0.25% glutaraldehyde for post-fixation (protocol variant 12 in Suppl. Table 1). Additionally, we omitted the denaturation step, since it may have a negative impact on the chromatin structure. However, the tested fixation conditions did not improve the preservation of the chromatin structure. Moreover, we observed nuclei less expanded (∼47 µm; (Suppl. Fig. 4) than nuclei fixed in a FA/AA mixture (∼61 µm; Fig. 6h). Thus, the applied fixation solutions influence the ability of the nuclei to expand, and the fixation in a mixture of 1% FA and 1% AA is most suitable for ExM.

The expansion of nuclei is a physical process that might alter the native structure of chromatin if nuclei were strongly fixed. Therefore, we tested whether a gentle fixation of root tips in 1% FA + 1% AA for 20 min could improve the preservation of the chromatin structure. Two different monomer solutions were applied, original MS1, and MS2 with less acrylamide concentration (Suppl. Table 2). In the case of MS2, we performed enzymatic digestion of nuclei using proteinase K (protocol variants 17-20 in Suppl. Table 1). In both cases, expanded nuclei were observed, but the chromatin structure was better preserved by MS2 (Suppl. Fig. 5). In addition to chromatin also nucleoli were strongly labelled by DAPI suggesting that the proteinase K treatment has an impact on the stability of chromatin. Hence, we omitted the digestion with proteinase K from the protocol (Suppl. Table 1 protocol variant 21). Thus, we obtained expanded nuclei with chromatin structures already visible by WF, but with nucleoli free of DAPI labelling (Figs. 4 a, d, g). SIM, DCV and WF showed comparable images. Further magnification revealed network-like structures in all three imaging methods (Figs. 4b, c, e, f, h, i). The protocols delivering less expanded nuclei or very little preserved chromatin were omitted from the further optimization process. The protocol (1% FA/AA in MS1 or MS2, no denaturation/digestion) showing the best-expanded nuclei was used in three independent experiments.

### Except for their ultrastructure, the global chromatin arrangement of expanded nuclei is preserved after ND-FISH and immunolabelling

Expansion microscopy in combination with ND-FISH for the visualization of high-copy repeats in expanded nuclei was tested next. A DNA denaturation step, an essential part of the standard FISH procedure was omitted since it was shown that the detection of barley centromere and telomere repeats does not require this step (**Cuadrado et al. 2009**). Using a barley centromere-specific probe we detected by WF microscopy in expanded nuclei hybridization signals corresponding to centromeres arranged in Rabl orientation, which originates from the former arrangement of the centromeres during telophase (**Rabl 1885**), an interphase organization also common for barley (**Schubert et al. 2016**) (Fig. 5). However, SIM revealed that the arrangement of chromatin in fibres and domains was damaged (Fig. 5j, k, l) l. Thus, the position of repeats after nuclei expansion was maintained but the chromatin ultrastructure organized in a network-like manner was lost. The overall quality of the chromatin structure was low compared to the protocol without ND-FISH, presumably due to the additional treatment steps required for ND-FISH. The overnight incubation in 50% deionized formamide could have a negative impact on the preservation of chromatin.

To investigate whether proteins can be visualized in expanded barley nuclei we employed indirect immunostaining using antibodies against the centromere-specific histone variant CENH3 of barely (**Houben et al. 2007**). In contrast to FISH, instead of monomer solution MS2, we used solution MS1. Since indirect immunostaining requires several incubation and re-expansion steps and MS2 delivers fragile gels due to the low acrylamide amount it is challenging to conduct the protocol without damaging the gel. The immunofluorescence signals of CENH3 could be detected independently of the used monomer solution. Prior to expansion, no denaturation/digestion was carried out.

Similar to the centromere-specific repeats identified by FISH the CENH3 signals displayed the expected Rabl-like positions in expanded and unexpanded nuclei (Fig. 6). Imaged by WF the immunosignals were homogeneous. But SIM revealed that the CENH3 signals were scattered in expanded nuclei (Fig. 6j). In contrast, the CENH3 and overall chromatin structures of expanded nuclei were well preserved (Fig. 6 d-f).

In summary, ExM allows enlarging isolated plant nuclei physically with an expansion factor of ∼4.2-times. Barley nuclei with a size between ∼40-110 µm were obtained. Distinct chromatin domains can be detected in expanded nuclei without the need for optical super-resolution microscopy (Fig. 4). Barley centromeres can be visualized by combining ExM with ND-FISH or indirect immunostaining. However, super-resolution microscopy revealed that ExM results in the impairment of the network-like organization of the chromatin substructure.

## Discussion

We adapted expansion microscopy (ExM) protocol for isolated plant nuclei. With the aim to preserve the chromatin structure we adapted the U-ExM protocol of **Gambarotto et al. (2019**) which was initially established for centrioles of *Chlamydomonas*. To ensure the best possible chromatin structure maintenance we tested different fixatives such as formaldehyde, glutaraldehyde and glyoxal. It is known that fixation has a crucial effect on chromatin structure preservation (**Kozubek et al. 2000; Howat and Wilson 2014**). **Guillot et al. (2004)** demonstrated that fixation and cell permeabilization affects the distribution of RNA polymerase II molecules in human cells under conditions that do not sustain the cellular ultrastructure. While formaldehyde is routinely used for the fixation of specimens before immunolabelling and light microscopy (**Puchtler and Meloan 1985**), glutaraldehyde is commonly used for electron microscopy-based observations (**Hayat 1986; Park et al. 2016**). Glyoxal was successfully applied for different animal tissues to improve structural features and to reduce formaldehyde fixation artefacts (**Richter et al. 2018**). Fixation procedures required before applying immunostaining and FISH may induce structural artefacts within the specimens. However, **Markaki et al. (2012)** demonstrated that by appropriately adapted 3D-FISH the key characteristics of cell nuclei are preserved and that SIM discovers new insights into the functional nuclear organization.

Although all fixatives kept nuclear morphology and the nucleoli of barley nuclei, only formaldehyde allowed the expansion of ∼4.2-times. The mild fixation of roots using 1% FA and 1% AA with a 20 min incubation time provided better results than the stronger fixation of roots in 4% FA + 4% AA for 20 or 40 min. The sample preparation steps, denaturation and digestion, which are required to homogenize the mechanical properties of different non-plant tissues (**Chen et al. 2015; Cho et al. 2018; Wassie et al. 2019**) were omitted because both steps impaired strongly the chromatin structure. This observation is in agreement with previous studies showing that chromatin becomes damaged when isolated nuclei were exposed to a detergent (**Szabó et al. 1990**). A similar negative effect has been described for proteinase K that cleaves chromatin into 50 kb fragments (**Szabó et al. 1990; Gal et al. 2000**). We speculate that this might be the reason that DAPI-specific signals were found within the nucleolus after proteinase K treatment. Further, DNA damage leads to the accumulation of RNAs and proteins inside of the nucleolus (**Lindström and Latonen 2013; Jin et al. 2014**). Moreover, **Kao and Nodine (2019)** showed that a mild proteinase K treatment impairs the fluorescence signal intensities after immunolabelling in expanded *Arabidopsis* ovules and seeds.

ExM allowed us to visualize the main structural features of nuclei like nuclear shape and the nucleoli and to observe chromatin structures invisible in unexpanded nuclei when detected with classical WF. To further increase the resolution of expanded specimens and to check the substructure maintenance of chromatin we applied super-resolution microscopy after ExM. We identified network-like organized chromatin, similar to that observed in unexpanded nuclei of mammals (**Markaki et al. 2012**) and plants (**Ma et al 2017; Schubert 2017**).

However, the combination of ExM with SIM did not result in more structural information, although the achieved SIM resolution (∼25-35 nm) in expanded nuclei was higher than in unexpanded ones (∼50-60 nm). Instead, the visualization of chromatin in unexpanded nuclei by SIM delivers better results (compare Figures 2 and 4). What could be the reason? **Pernal et al. (2019)** showed that expansion is anisotropic not only between different tissues but also between different subcellular compartments and even within subcellular compartments themselves. This observation may be a reason that the chromatin ultrastructure within nuclei becomes damaged and is difficult to preserve even after appropriate fixation. The employed 63×/1.4 oil objective (working distance of 0.19 mm) required for SIM, caused another technical challenge due to the impossibility to analyse nuclei which were distantly embedded from the coverslip. To circumvent this problem, **Cahoon et al. (2017)** prepared cryo-sections and successfully observed the expanded synaptonemal complex of the fruit fly. But this approach is laborious and not suitable for high-throughput experiments.

We combined ExM with ND-FISH to detect centromeric repeats in barley. The position of the detected fluorescence signals corresponded to centromeric signals observed in original, unexpanded nuclei. Thus, a nucleus that underwent expansion maintains its general morphology and chromatin organization. On the other hand, SIM revealed that the ultrastructure of centromeric chromatin was only partially preserved after ExM and ND-FISH. Moreover, chromatin containing DNA molecules creating a network-like structure (**Beseda et al. 2020)** is more complex than short mRNA molecules. The combination of ExM and FISH was successfully used to visualize mRNA in expanded mammalian cell cultures and brain tissue (**Chen et al. 2016; Wang et al. 2018a**), but their protocols differ from our protocols by using different fixation solution and omitting the denaturation/digestion steps. Moreover, compared to the relatively short mRNA molecules, chromatin is more complex by creating network-like structures in animals and plants (**Markaki et al. 2012; Schubert 2014; Beseda et al. 2020**), and thus can collapse more easily during the process of physical magnification. Therefore, we speculate that current ExM protocols can reveal and detect the correct RNA positions using FISH, but sustaining the chromatin ultrastructure is more challenging. Similar to the FISH experiments we localized the centromere-specific protein CENH3 in the correct Rabl orientation, but again the chromatin ultrastructure was impaired. This observation is reasonable because CENH3, as a component of nucleosomes, is associated with DNA forming centromeric heterochromatin.

On the other hand, several reports (**Chozinski et al. 2016; Cahoon et al. 2017; Freifiled et al. 217; Jiang et al. 2018; Wang et al. 2018b; Gambarotto et al. 2019; Kao and Nodine 2019; Xu et al. 2019)** demonstrated the improved visualisation of target proteins after applying ExM. Thus, ExM can reveal the unaltered localization of target molecules, but preserving the chromatin ultrastructure of isolated nuclei is more challenging, and therefore further improved ExM protocols have to be developed. Otherwise, expanded chromatin structures imaged by wide-field microscopy will not deliver more information as achieved by super-resolution microscopy on unexpanded structures.

## Abbreviations

3D: 3-dimensional
AA: acrylamide
APS: ammonium persulfate
DAPI: 4′,6-diamidino-2-phenylindole
DCV: deconvolution
FA: formaldehyde
FISH: fluorescence in situ hybridization
GA: glutaraldehyde
h: hour
min: minutes
MS: monomer solution
PBS: phosphate buffer saline
PLL: poly-L-lysine
RT: room temperature
SIM: structured illumination microscopy
TEMED: tetramethylethylenediamine
U-ExM: ultrastructure expansion microscopy
WF: widefield

## Acknowledgements

This work has been supported by the Deutsche Forschungsgemeinschaft (Schu 762/11-1) and by the Czech Science Foundation (17-20613Y).

---

**Suppl. figure 1:**
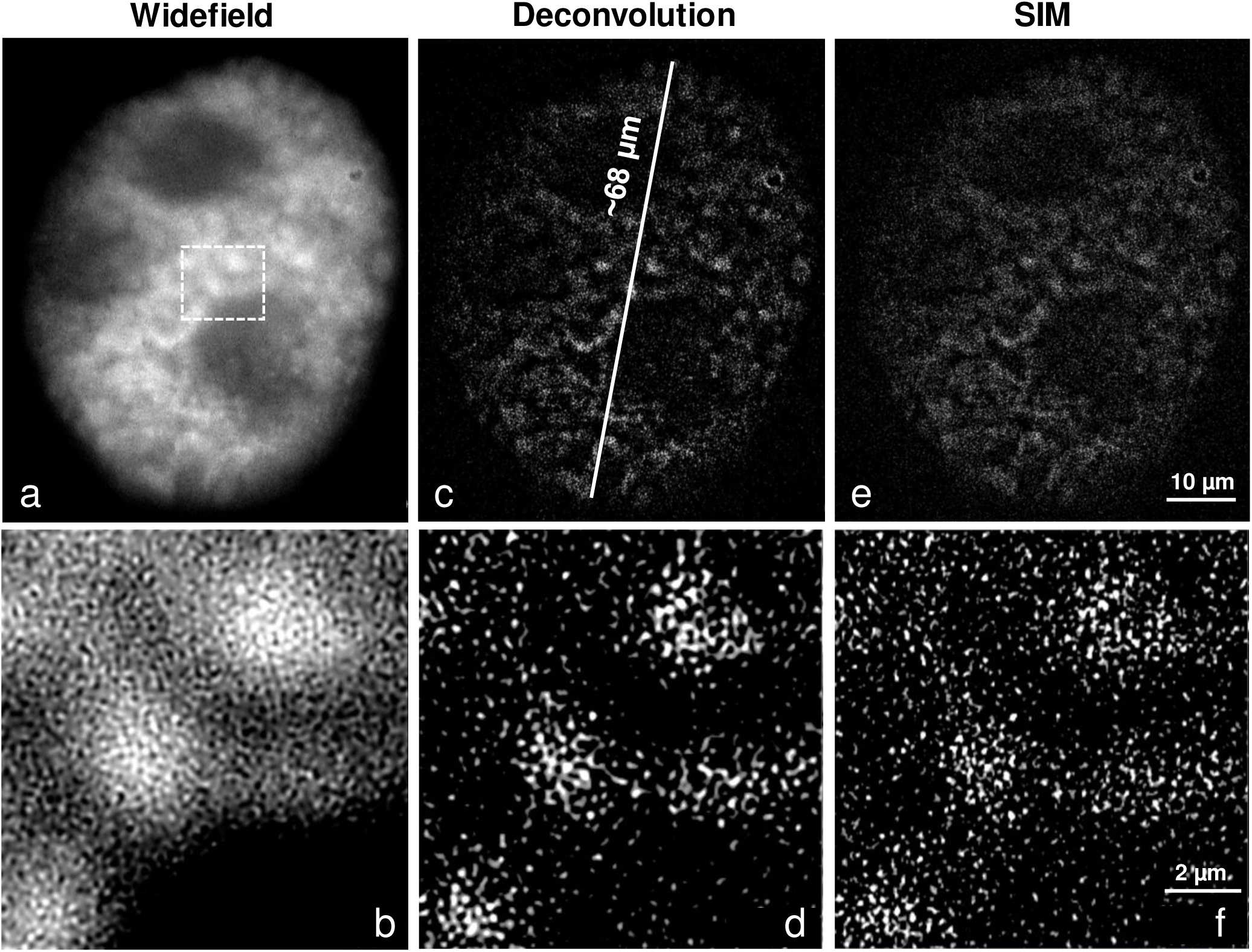
The chromatin fine structure is impaired by heat denaturation applying ExM protocol variant 2 (see suppl. table 1) to achieve complete expansion, as especially demonstrated by SIM of the magnified selected region (dashed rectangle). The expanded nucleus was imaged by WF (a, b), DCV (c, d) and SIM (e, f). Global chromatin was stained by DAPI. Global chromatin was stained by DAPI.

**Suppl. figure 2:**
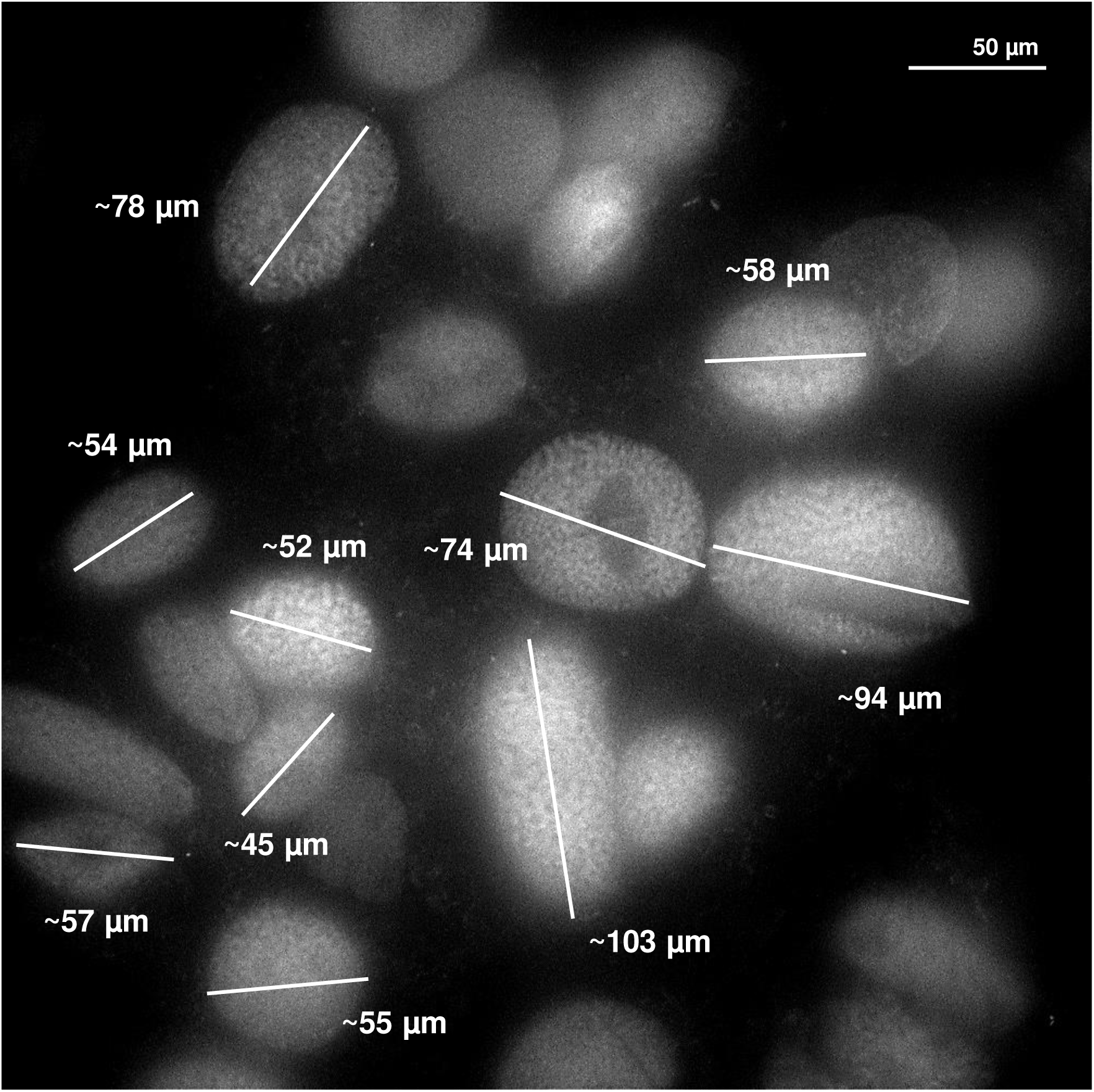
Mixture of different nuclei from root tips after expansion acquired by a 20×/0.8 objective. The nucleus population originates from different cell cycle stages (G1, S, G2) and tissue layers of the root tip. The differences in DNA content influence the size of the nuclei and thus a high size variability is also present after expansion.

**Suppl. figure 3:**
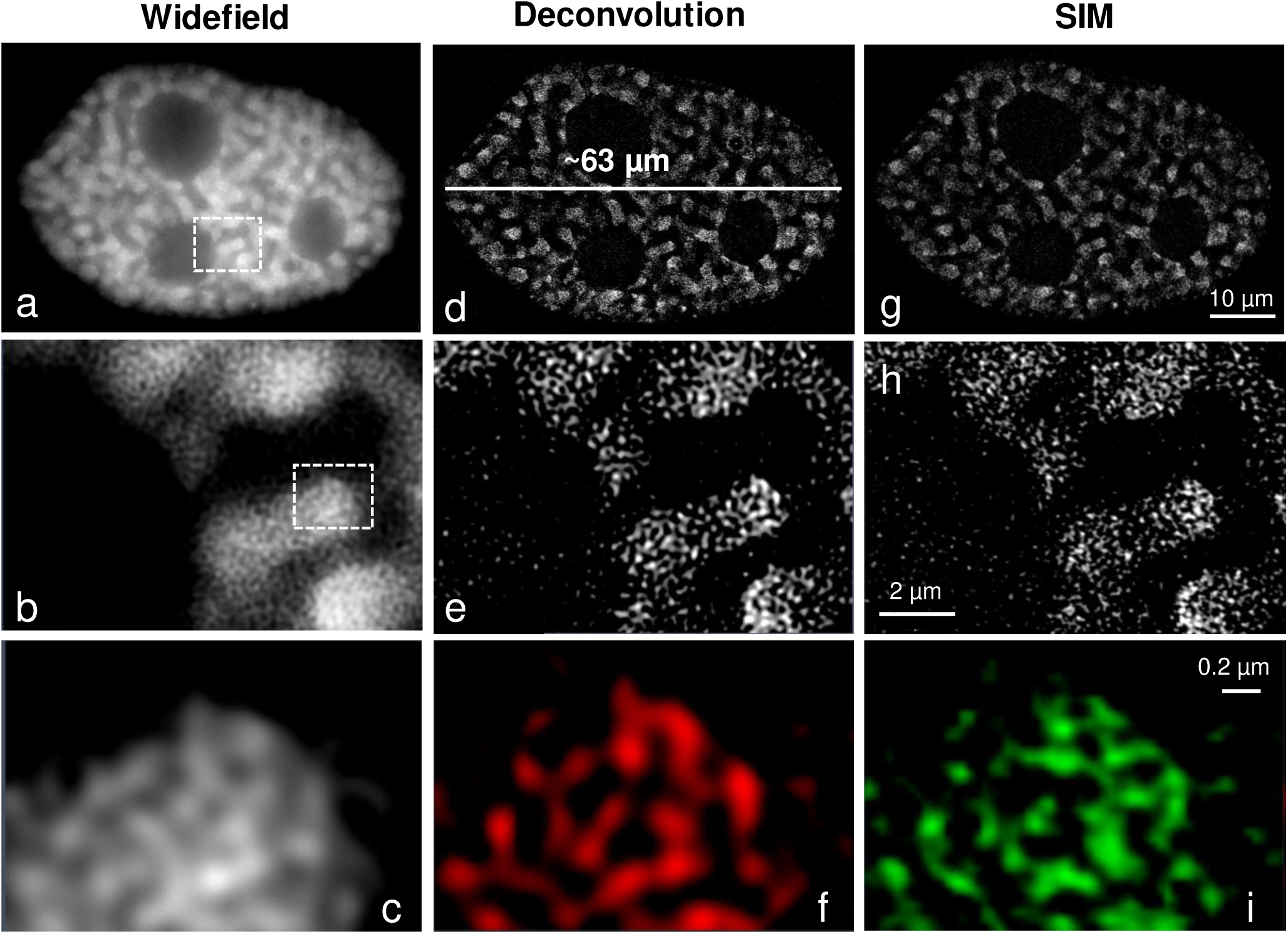
The chromatin fine structure is mainly preserved after denaturation with NaOH applying ExM protocol variant 3 (see suppl. table 1) to achieve complete expansion. By WF (a, b, c), DCV (d, e, f) and SIM (g, h, i) identical chromatin fine structures labelled by DAPI become visible as especially demonstrated at the highest magnification (bottom panel) of the magnified selected regions (dashed rectangles).

**Suppl. figure 4:**
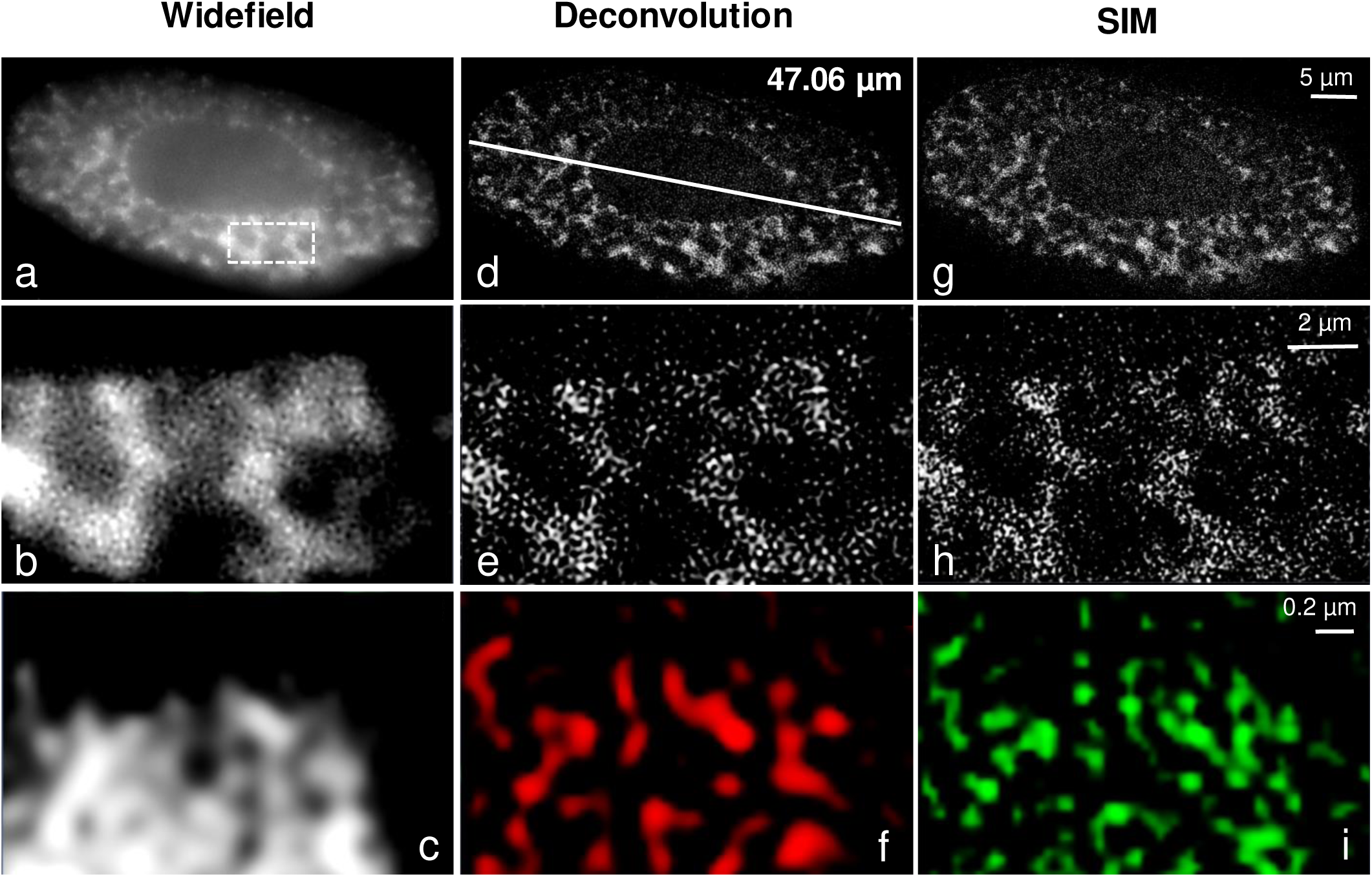
The fixation with FA/GA and post-fixation with GA (ExM protocol variant 12, see suppl. table 1) delivered less expanded nuclei than the FA/AA fixation. The chromatin domains visible in WF (a-c). DCV (d-f) and SIM (g-i) showed that the native network-like chromatin structure was impaired.

**Suppl. figure 5:**
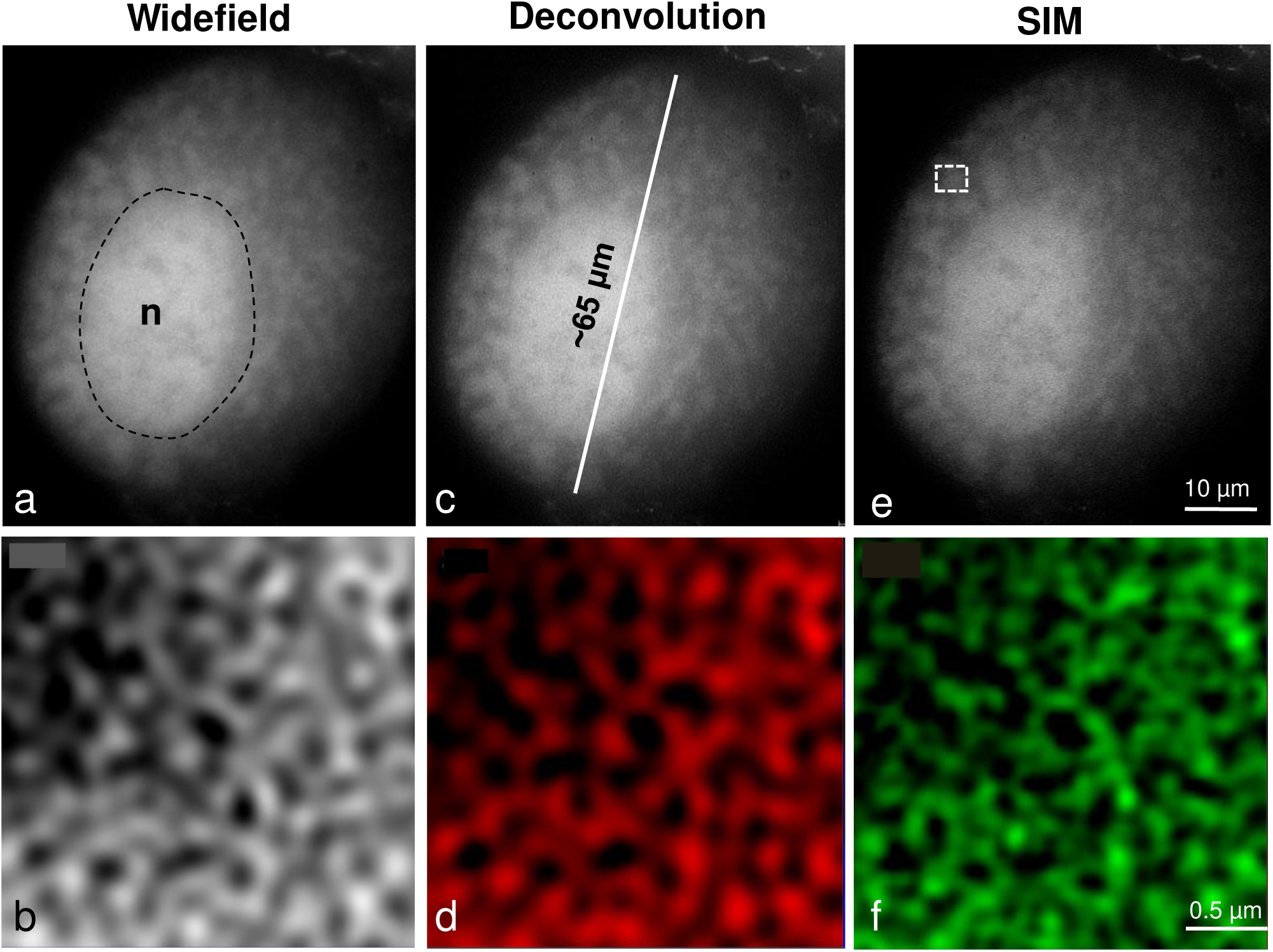
Although proteinase K treatment (protocol variant 18 in suppl. table 1) does not impair the chromatin fine structure the digestion induces unspecific nucleolus (n) labelling. By WF (a, b), DCV (c, d) and SIM (e, f) imaging, especially at high magnification (bottom panel) of the selected region (dashed rectangle) similar chromatin structures labelled by DAPI become visible.

**Suppl. table 1:**
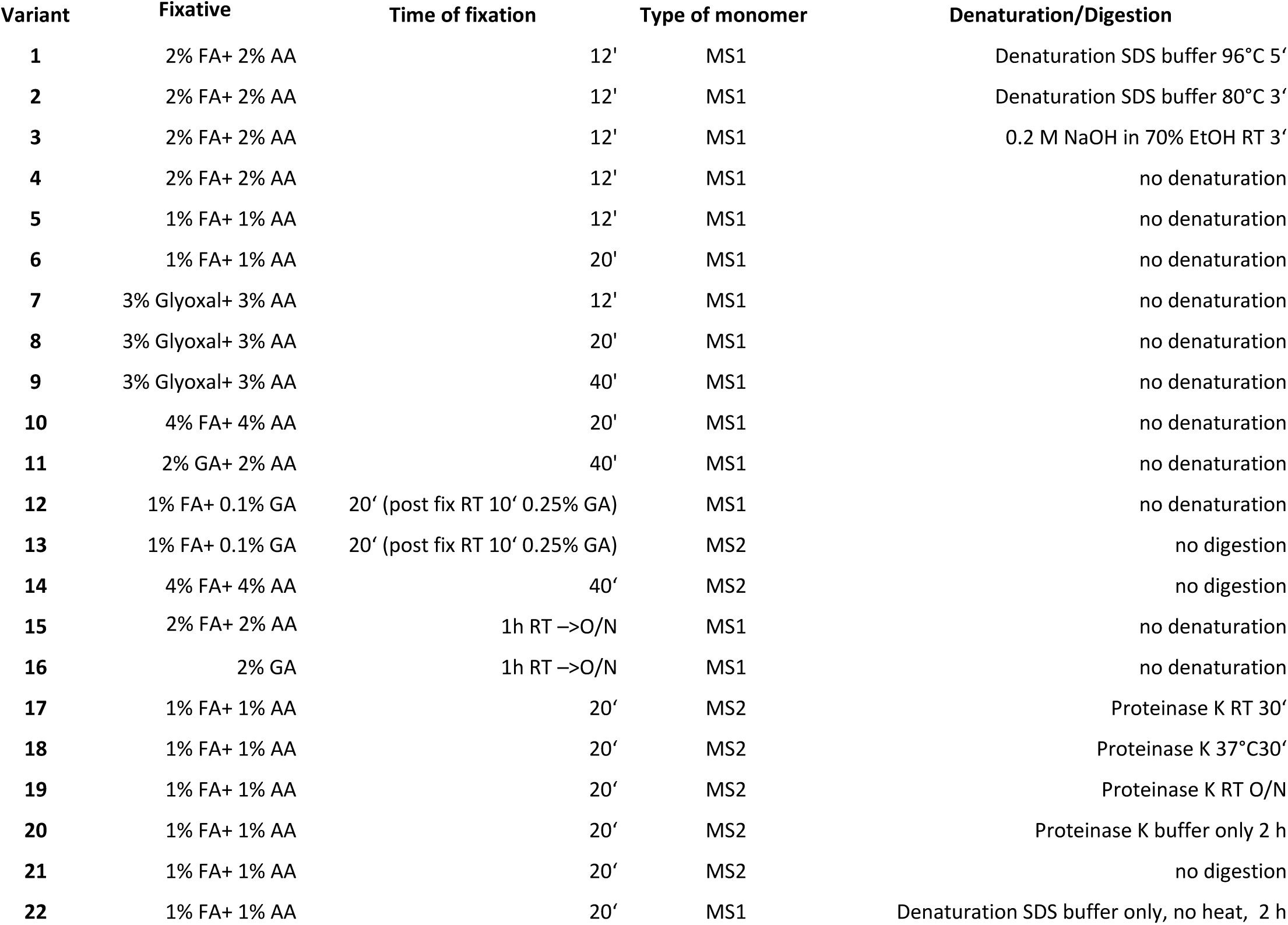
Protocol variants for plant nucleus ExM.

| Variant | Fixative | Time of fixation | Type of monomer | Denaturation/Digestion |
| --- | --- | --- | --- | --- |
| 1 | 2% FA+ 2% AA | 12' | MS1 | Denaturation SDS buffer 96°C 5' |
| 2 | 2% FA+ 2% AA | 12' | MS1 | Denaturation SDS buffer 80°C 3' |
| 3 | 2% FA+ 2% AA | 12' | MS1 | 0.2 M NaOH in 70% EtOH RT 3' |
| 4 | 2% FA+ 2% AA | 12' | MS1 | no denaturation |
| 5 | 1% FA+ 1% AA | 12' | MS1 | no denaturation |
| 6 | 1% FA+ 1% AA | 20' | MS1 | no denaturation |
| 7 | 3% Glyoxal+ 3% AA | 12' | MS1 | no denaturation |
| 8 | 3% Glyoxal+ 3% AA | 20' | MS1 | no denaturation |
| 9 | 3% Glyoxal+ 3% AA | 40' | MS1 | no denaturation |
| 10 | 4% FA+ 4% AA | 20' | MS1 | no denaturation |
| 11 | 2% GA+ 2% AA | 40' | MS1 | no denaturation |
| 12 | 1% FA+ 0.1% GA | 20' (post fix RT 10' 0.25% GA) | MS1 | no denaturation |
| 13 | 1% FA+ 0.1% GA | 20' (post fix RT 10' 0.25% GA) | MS2 | no digestion |
| 14 | 4% FA+ 4% AA | 40' | MS2 | no digestion |
| 15 | 2% FA+ 2% AA | 1h RT →O/N | MS1 | no denaturation |
| 16 | 2% GA | 1h RT →O/N | MS1 | no denaturation |
| 17 | 1% FA+ 1% AA | 20' | MS2 | Proteinase K RT 30' |
| 18 | 1% FA+ 1% AA | 20' | MS2 | Proteinase K 37°C30' |
| 19 | 1% FA+ 1% AA | 20' | MS2 | Proteinase K RT O/N |
| 20 | 1% FA+ 1% AA | 20' | MS2 | Proteinase K buffer only 2 h |
| 21 | 1% FA+ 1% AA | 20' | MS2 | no digestion |
| 22 | 1% FA+ 1% AA | 20' | MS1 | Denaturation SDS buffer only, no heat, 2 h |

**Suppl. table 2:** Monomer and buffers solutions.

| **Monomer solution 1 (MS1)** | **Final concentration** |
| --- | --- |
| Sodium acrylate 38% | 19% |
| Acrylamide 40% | 10% |
| N,N'-methylenebisacrylamide 1% | 0.1% |
| PBS 10× | 10× |
| ddH<sub>2</sub>O | Fill up to final volume |
| TEMED 10% | 0.2% |
| APS 10% | 0.2% |

| **Monomer solution 2 (MS2)** |  |
| --- | --- |
| Sodium acrylate 38% | 8.6% |
| Acrylamide 40% | 2.6% |
| N,N'-methylenebisacrylamide 1% | 0.15% |
| NaCl | 2 M |
| PBS 10× | 1× |
| ddH<sub>2</sub>O | Fill up to final volume |
| TEMED 10% | 0.2% |
| APS 10% | 0.2% |

| **Digestion buffer** |  |
| --- | --- |
| Tris-Base | 50 mM |
| EDTA | 1 mM |
| Triton X-100 | 0.5% |
| guanidine HCl | 0.8 M |
| ddH<sub>2</sub>O | Fill up to final volume |
| add freshly proteinase K | 8 U/ml |

| **Denaturation buffer** |  |
| --- | --- |
| Sodium dodecyl sulphate (SDS) | 200 mM |
| NaCl | 200 mM |
| Tris-Base | 50 mM |
| ddH<sub>2</sub>O | Fill up to final volume |

